# Community complexity does not weaken pairwise coevolution in a soil bacterial community

**DOI:** 10.1101/2025.07.01.662545

**Authors:** Zoltan Erdos, Daniel Padfield, Elze Hesse, Angus Buckling, Meaghan Castledine

## Abstract

Exploitative interactions, such as predator-prey and host-parasite interactions, are ubiquitous in microbial communities. These interactions shape community density and composition, imposing strong selection on members to evolve countermeasures reducing the negative impacts of exploitation. Exploitative coevolution is often studied between species pairs in isolation, which may over-estimate the strength and relevance of pairwise coevolution. Here we studied how community context influences coevolution between *Pseudomonas fluorescens* (exploited) and *Variovorax sp*. (exploiter). We evolved these species in pairwise coculture and embedded within a five-species community to investigate evolved changes in pairwise interactions. We found evidence for asymmetrical coevolution: *Variovorax* evolved more rapidly than *Pseudomonas*, leading to increased exploitation through time, while *Pseudomonas* evolved increased tolerance to *Variovorax* with time lag. The pairwise coevolutionary dynamics were not affected by the presence of other community members. Understanding how coevolutionary patterns change with increasing community complexity can have important implications for community persistence and function.

## Introduction

Exploitative interactions - where one species benefits at the expense of another – can result in the adaptive evolution of defence, counter-defence or reciprocal evolution of these traits (antagonistic coevolution) (Brockhurst & Koskella, 2013; Gandon et al., 2008). Antagonistic coevolution can have far-reaching consequences for ecological and evolutionary dynamics of communities; particularly so in microbial communities where organisms often have large populations and short generation times (Barraclough, 2015; Garbutt et al., 2011; Paterson et al., 2010), meaning ecological and evolutionary processes often happen simultaneously (Loreau et al., 2023). Previous work into antagonistic coevolution has primarily focussed on interactions between trophic levels, hosts and parasites (Friman & Buckling, 2013; Gómez & Buckling, 2011) and predator and prey (Friman et al., 2011; Johnke et al., 2017). However, it is also important within trophic levels where interactions often occur over exploitation of extracellular compounds (e.g. evolution of resistance to antibiotics via competition (Koch et al., 2014), evolution of increased competitiveness and exploitation in biofilms (Hansen et al., 2007; Kim et al., 2014)).

Community context may have a significant effect on pairwise coevolution (Barraclough, 2015; Blazanin & Turner, 2021; Manriquez et al., 2021). Being embedded within a community is likely to reduce the occurrence and speed of pairwise coevolution. Interacting with multiple community members will potentially reduce the frequency of interaction for a given species pair, result in trade-offs between adaptation to multiple species (Alseth et al., 2019; Friman & Buckling, 2013)) and increase the magnitude of trade-offs between abiotic and biotic adaptation (Briscoe Runquist et al., 2020; Gómez & Buckling, 2013; Hall et al., 2018; Lawrence et al., 2012; Luján et al., 2022; Yin et al., 2023), all of which will reduce the strength of reciprocal selection. Furthermore, reduced population sizes with increasing community members will reduce the supply of mutation on which selection acts (Castledine et al., 2020; Hart et al., 2019). Our understanding of how community complexity affects coevolutionary dynamics is however primarily limited to studies of bacteria-virus (bacteriophage) systems.

Here, we employ a time shift (Buckling & Rainey, 2002; Gaba & Ebert, 2009) approach to quantify the impact of community complexity on the coevolutionary dynamics of two focal species that engage in an exploitative interaction. *Variovorax* sp. (AB1) benefits from unknown metabolites produced by *Pseudomonas fluorescens* (AB1) (hereafter referred to as *Variovorax* and *Pseudomonas*), while the growth of *Pseudomonas* is reduced by the presence of *Variovorax* suggesting the potential for strong selection for coevolution (Castledine et al., 2024). This soil microbial community consists of five species (including *Pseudomonas* and *Variovorax*) and is dominated by competitive interactions (Castledine et al., 2024). Briefly, under similar experimental conditions to our current work, *Pseudomonas* is negatively impacted by all other species. While *Variovorax* reduces the fitness of all other species, it experiences a growth benefit in the presence of three of the four other community members (with highest exploitation against *Pseudomonas*). Therefore, we expect that community context would reduce the extent of pairwise coevolution between *Variovorax* and *Pseudomonas* by decreasing mutation supply and the strength of reciprocal selection. We hypothesize that strong selection would result in *Pseudomonas* and *Variovorax* evolving via arms race dynamics (e.g. *Pseudomonas* getting better at defence against exploitation and *Variovorax* evolving increased exploitation) in coculture, and that coevolution of the focal species would be weakened in a multispecies community.

## Materials and methods

### Experimental evolution

Experimental evolution treatments were set up to study adaptation and pairwise coevolution between *Pseudomonas* and *Variovorax* in different biotic conditions (Figure 1). A monoculture evolution treatment was set up to isolate any effects of correlated abiotic adaptation, which can alter competitive hierarchies, mediate exploitative dynamics, and influence niche partitioning within the community. A coculture and community evolution treatment was used to study coevolution in a pairwise and community background respectively, and to assess the impact of community complexity on pairwise coevolution between two species.

**Figure 1.**
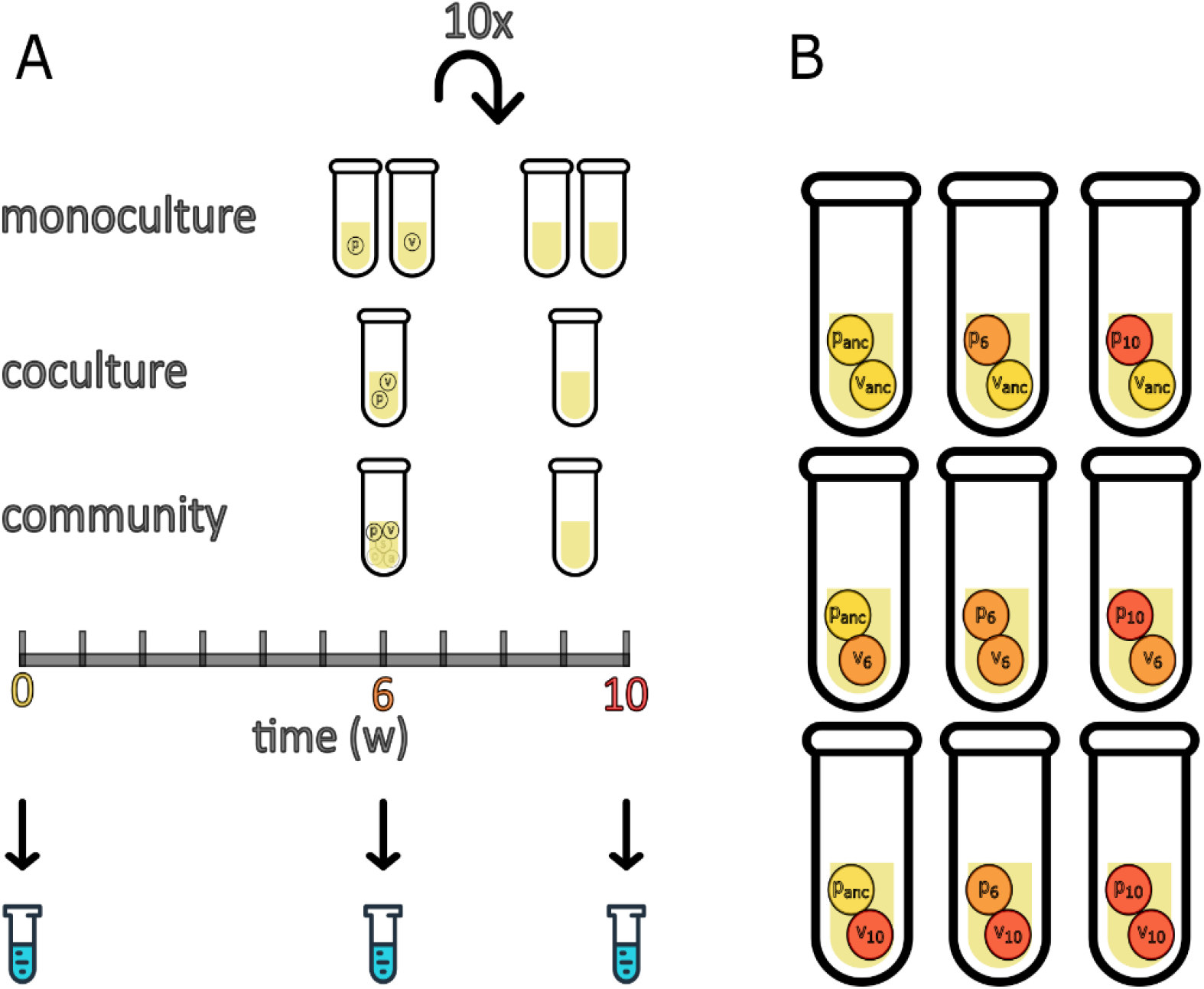
Experimental design to test the impact of community complexity on coevolution between focal species Pseudomonas (P) and Variovorax (V). The evolution experiment consisted of three treatments: monoculture, coculture and community. Each treatment has been passaged (1%) weekly for 10 weeks (A). Coevolution was assessed by growth assays combining Pseudomonas and Variovorax from different evolutionary timepoints from the same replicate line within each treatment (B). anc = ancestor, 6 = 6-week isolates, 10 = 10-week isolates.

The community evolution treatment had been carried out previously and results have been published in Castledine et al. (2020). We randomly selected eight replicate communities in our current study. Mono- and coculture evolution treatments were initiated with eight replicates (eight replicates for each species per monoculture) using the same *Pseudomonas* and *Variovorax* ancestors and culturing protocol. Briefly, bacterial isolates obtained from soil samples, including *Achromobacter sp*., *Ochrobactrum sp*., *Pseudomonas sp*., *Stenotrophomonas sp*., and *Variovorax sp*., were identified based on their distinct colony morphologies on King’s medium B (KB) agar. Each species was cultivated from a single colony in isolation for two days in 6 ml 1/64 Tryptic Soy Broth (TSB) medium at 28°C in glass microcosms. Inoculated abundances (colony forming unit, CFU) of each species were estimated approximately from optical densities (OD_600_; wavelength 600 nm) after two days of growth (equations for converting OD_600_ to CFU/μL described previously (Castledine et al., 2024) and adjusted to: 2×10^6^ CFU/µl. Replicate lines of communities (all species combined), cocultures (*Pseudomonas* and *Variovorax*) and monocultures (*Pseudomonas, Variovorax*) were established using a 20 μL inoculum from each species into fresh 6 mL 1/64 TSB. Cultures underwent weekly serial 100-fold dilutions (transfer of 1% inocula into fresh media) over ten weeks, with samples frozen every second transfer (−70 °C in glycerol, final concentration: 25%). Frozen samples from the ancestor (used as inoculum above), 6- and 10-week-old cultures were plated onto KB agar and incubated for 2 days at 28 °C. Six clones of *Pseudomonas* and *Variovorax* each per community, coculture and monoculture evolution line (replicate) were isolated from each timepoint and grown for 48 hours in 1/64 TSB before being combined and frozen at −70 °C in glycerol to be used in time-shift assays.

### Time-shift assays

To test whether the two focal species, *Pseudomonas* and *Variovorax*, have coevolved over the 10-week period we conducted time-shift assays, where one species was cultured with the other from a population of the past, contemporary or future timepoints. These assays were carried out in the absence of the rest of the community. This approach allows for signatures of different types of coevolution to be detected. Arms race dynamics (ARD, i.e. selection of defence and counter defence) is directional, with focal species having a greater abundance when competing with the other species isolated from a past time point, and lower densities when competing against competitors from a future time point. If instead selection on specific genotypes fluctuates through time (fluctuating selection dynamic, FSD), performance may be particularly high or low for contemporaneous interactions (Hall et al., 2011).

Populations of *Pseudomonas* and *Variovorax* from one of the three different timepoints of the evolution experiment (ancestor, 6 week and 10 week) were reassembled, resulting in 9 pairwise combinations per treatment (Figure 1B). Cultures of *Pseudomonas* and *Variovorax* were assembled using all 6 clones from the same treatment replicate. Eight replicate microcosms were set up for each combination within the three different evolution backgrounds except for monocultures, see below. The culture conditions of all treatments were established as described for the initial experimental evolution experiment, with approximately 2×10^6^ CFUs per species inoculated into fresh microcosms. After one week, culture samples were cryogenically frozen and then plated onto KB agar. Population densities were estimated by counting the number of CFUs (10^5^ diluted) after two days of growth at 28 °C. We use proportion of *Pseudomonas* to interpret coevolutionary dynamics. Proportion of species is often more insightful than densities in describing coevolution as it highlights shifts in relative abundance of each species. These shifts are indicative of selective pressure imposed by interspecies interaction (e.g. parasitism) and is less affected by fluctuations in total population size that can be influenced by various factors such as stochastic variation in density between microcosms.

Three replicates in the monoculture evolution line (for both *Pseudomonas* and *Variovorax*) were contaminated prior to week 6, therefore monoculture growth assays were carried out with only 5 replicates. Three replicates of the community evolution line were contaminated during the time-shift assay, and these were removed from the analysis. The total density of pairwise combinations were assessed using the total abundance (combined CFU) of *Pseudomonas* and *Variovorax* in each microcosm.

### Statistical analyses

All data analyses were carried out in R v 4.3.1 (R Core Team, 2025). First, we analysed whether the density of each species varied in their interaction with contemporary counterparts over time. For this model, the time-shift assays were subset to retain only the bacterial densities from their contemporary assays. The density of each species was analysed in linear mixed-effects models (LMM) separately, with log_10_ transformed density as the response variable, treatment, time and their interactions as explanatory variable and random intercepts fitted for each replicate to account for non-independence of observations.

One replicate of the time-shift assays in the coculture treatment (*Pseudomonas* time 6, *Variovorax* time 10) showed an unusually high *Variovorax* count and was found to be an influential outlier (Grubbs-test) and was removed from the analysis. To test how community complexity affects coevolutionary dynamics we used a binomial generalized linear mixed-effects models (GLMMs) with a logit link function, the proportion of *Pseudomonas* as combined binary response variable and treatment (complexity) x evolutionary time of *Pseudomonas* x evolutionary time of *Variovorax* fitted as fixed effects, as well as their 3-way interaction. We included random intercepts for each replicate line to account for non-independence of observations. To study coevolution within treatments, we tested the effect of coevolutionary time of both *Pseudomonas* and *Variovorax*, plus their interaction, on the proportion of *Pseudomonas* in separate models per treatment using binomial GLMMs with a logit link function for each evolutionary background. Total density of pairwise combinations were analysed with a linear mixed-effects model (LMM) with log_10_ transformed total density as the response variable, treatment, evolutionary time of *Pseudomonas* and *Variovorax* and their interactions as explanatory variables and random intercepts fitted for each replicate to account for non-independence of observations. Time was included as discrete variable in all analysis. These analyses employed LMMs and GLMMs using the ‘*lme4’* package (Bates et al., 2015). For these LMMs and GLMMs, we used the ‘*DHARMa’* package (Hartig, 2018) to check residual behaviour and model simplification was carried out using likelihood ratio test. Post-hoc multiple comparison tests on the most parsimonious models were carried out using the R package ‘*emmeans’* (Lenth, 2023), using Tukey adjustment.

## Results

### Population densities change across contemporary species pairs evolved within community

We first determined how the relative success of *Variovorax* and *Pseudomonas* differed between treatments and through time. Specifically, we determined the density of each species when cultured with their contemporary counterparts. The effect of treatment (monoculture, co-culture, 5 species community) on *Pseudomonas* density did not differ through time (LMM: community complexity x evolutionary time: χ2_2_ = 3.06, p = 0.22, Figure 2), nor was there an overall effect of time (LMM: evolutionary time: χ2_1_ = 3.14, p = 0.077). Treatment did however have a significant effect on *Pseudomonas* density (LMM: community complexity: χ2_3_ = 21.20, p < 0.001, Figure 2). *Pseudomonas* populations that had evolved in community (4.52 log10 CFU/ml [4.43, 4.60]; Tukey HSD: estimate = 0.21, t-ratio = 3.1, p = 0.018) reached significantly lower densities compared to the ancestor and those evolved as cocultures (Tukey HSD: estimate = 0.14, t-ratio = 2.77, p = 0.051). Mean *Pseudomonas* population density in contemporary combinations of monoculture evolved *Pseudomonas* and *Variovorax* (4.79 log_10_ CFU/ml; 95% CI: [4.69, 4.88]) did not differ significantly from that of the ancestor *Pseudomonas* and ancestor *Variovorax* combination (4.73 log_10_ CFU/ml; 95% CI: [4.62, 4.84]; Tukey HSD: estimate = -0.057, t-ratio = -0.80, p = 0.85). Similarly, mean *Pseudomonas* density in contemporary combinations of coculture evolved lineages (4.65 log_10_ CFU/ml; 95% CI: [4.57, 4.73]; Tukey HSD: estimate = 0.078, t-ratio = 1.19, p = 0.64) were not different to the ancestor combination. Evolving in a community, and coculture to a lesser extent, results in lower *Pseudomonas* densities when cultured with *Variovorax*, suggesting a change in their interaction compared to the ancestors or monoculture evolved lineages.

**Figure 2.**
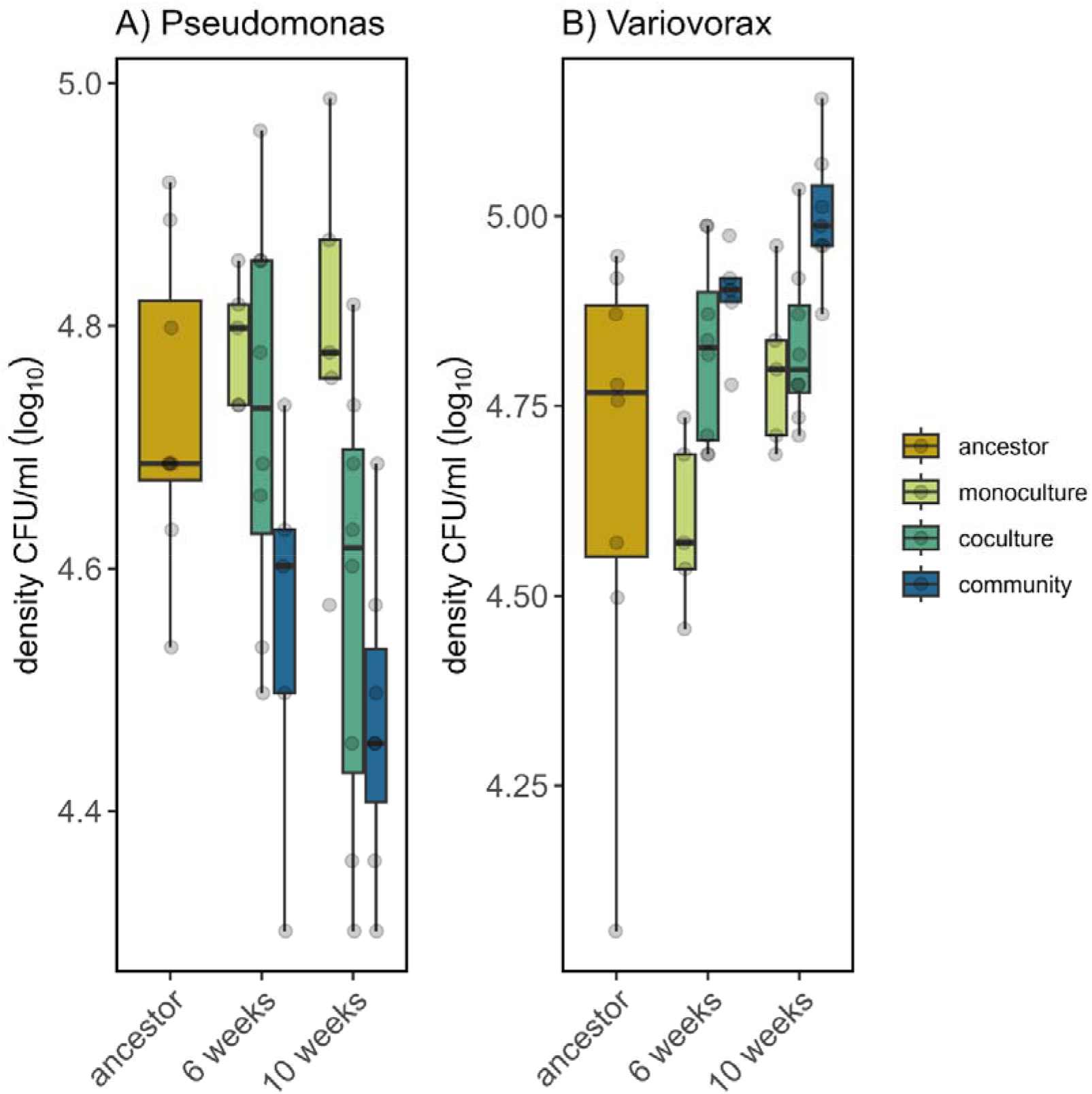
Density of Pseudomonas (A) and Variovorax (B) in their contemporary combinations of ancestor and evolved lineages in monoculture, coculture or community culture. Population densities are plotted against evolutionary time. Boxplots showing medians, first and third quartiles, whiskers are 1.5 * interquartile range (IQR). Individual points represent individual treatment replicates.

Similarly to *Pseudomonas*, the density of *Variovorax* was unaffected by an interaction between treatment and evolutionary time (LMM: community complexity x evolutionary time: χ2_2_ = 4.29, p = 0.12, Figure 2) or evolutionary time itself (LMM: evolutionary time: χ2_1_ = 1.43, p = 0.23), but treatment had a significant effect on *Variovorax* density (LMM: community complexity: χ2_3_ = 21.14, p < 0.001, Figure 2). Mean contemporary population densities of *Variovorax* in the ancestor combination (4.67 log_10_ CFU/ml; 95% CI: [4.55, 4.80]) were not different to those evolved in monoculture (4.70 log_10_ CFU/ml; 95% CI: [4.60, 4.80]; Tukey HSD: estimate = -0.024, t-ratio = -0.30, p = 0.99) or coculture (4.83 log_10_ CFU/ml; 95% CI: [4.74, 4.92]; Tukey HSD: estimate = -0.15, t-ratio = -2.00, p = 0.21). However, mean *Variovorax* densities in the community evolution treatment reached significantly higher densities than the ancestor in their respective contemporary cocultures (4.95 log_10_ CFU/ml; 95% CI: [4.86, 5.05]; Tukey HSD: estimate = -0.28, t-ratio = -3.49, p = 0.007). These results confirm that community context has a significant impact on the adaptation of species, and that coevolution lead to a change in interaction between species (when compared to ancestor or monoculture evolved lines).

### Antagonistic coevolution evident between species pairs

Time shift assays were used to characterise coevolution in coculture and in a community by growing *Pseudomonas* with *Variovorax* isolated from different time-points and vice-versa (Figure 1). To control for abiotic adaptation, pairs of monoculture lines (*Pseudomonas* and *Variovorax* cultured alone) evolved alongside the coculture and community treatments were also subjected to time shift assays.

The results of the time shift assays are indicative of antagonistic coevolution between *Pseudomonas* and *Variovorax* in both coculture (GLMM: *Pseudomonas* time x *Variovorax* time: χ2_3_ = 7.15, p = 0.067, Figure 3B) and in the community (GLMM: *Pseudomonas* time x *Variovorax* time: χ2_3_ = 21.85, p < 0.001, Figure 3C). *Variovorax* showed a tendency to increase in competitiveness against *Pseudomonas* through time in both the coculture and community treatments. Compared to the ancestral *Variovorax*, populations evolved for either 6- or 10-weeks in both coculture (Figure 3B) and community (Figure 3C) treatments reduced the proportion of ancestral *Pseudomonas*. There was no significant difference between the effect of *Variovorax* coevolved for 6- or 10-weeks on the proportion of ancestral *Pseudomonas* for the coculture treatment (Tukey HSD_coculture_: estimate = -0.027, z-ratio = -0.15, p = 0.88), however ancestral *Pseudomonas* proportion was significantly higher in the community-evolved treatment, when cultured with 10-week *Variovorax* compared to 6-week *Variovorax* (Tukey HSD_community_: estimate = -0.50, z-ratio = -2.69, p = 0.007, Figure 3BC). This latter result could be attributed to effects associated with other species in the community evolution treatment (e.g. adaptation to other species).

**Figure 3.**
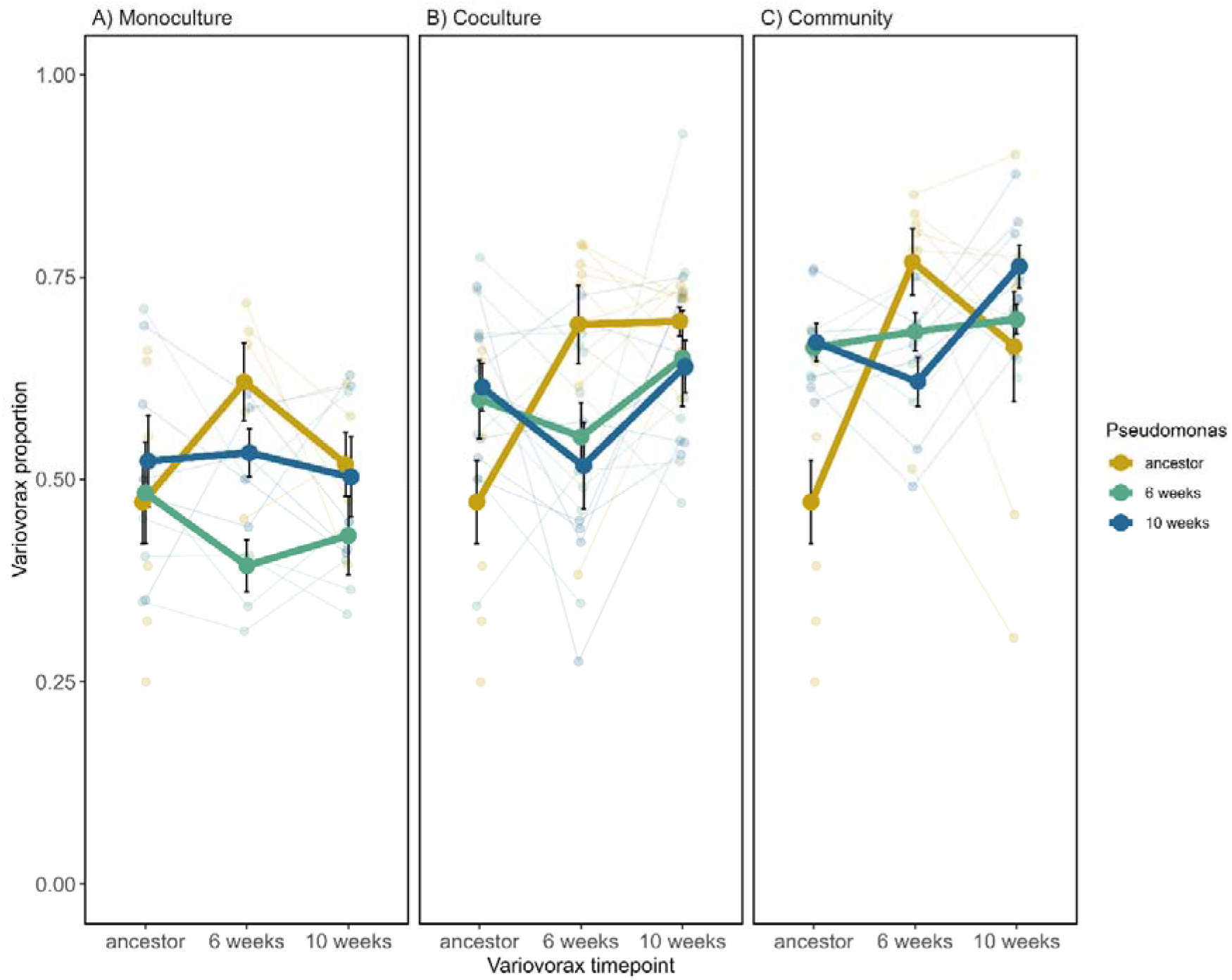
Relative proportion of Variovorax when cultured with Pseudomonas from different time-points. Pseudomonas and Variovorax evolved in (A) monoculture (each species in isolation), (B) in coculture (Pseudomonas and Variovorax) and in (C) community (Pseudomonas and Variovorax embedded with 3 other bacterial species)). We found a significant interaction between evolutionary time of Pseudomonas and Variovorax when species evolved as co-cultures (B) or within a community (C), but not when these had evolved in isolation (A). Independent treatment replicates are small points, large points are mean proportion of Variovorax, bars represent ± SE. Thin lines are connecting the replicates that are tracked through time (except for the ancestor), thick lines represent the mean. Ancestor-ancestor combinations between the panels are the same 8 replicates presented to aid visualization.

*Pseudomonas* underwent reciprocal adaptation to *Variovorax*. The proportion of *Pseudomonas* cultured with 6-week *Variovorax* increased with *Pseudomonas* evolutionary time for both coculture (Tukey HSD_ancestor – 6 week_: estimate = -0.66, z-ratio = -3.45, p = 0.0016; Tukey HSD_ancestor – 10 week_: estimate = -0.85, z-ratio = -4.37, p < 0.001, Figure 4B) and community (Tukey HSD_ancestor – 10 week_: estimate = -0.79, z-ratio = -3.07, p = 0.0061, Figure 4C) treatments. While *Pseudomonas* showed an overall increase in resistance through time against 6-week *Variovorax*, resistance declined against ancestral *Variovorax* for both treatments (Figure 4BC). Resistance to 10-week *Variovorax* was not different between any of the *Pseudomonas* evolutionary timepoints for either the coculture or community treatment. These dynamics are consistent with fluctuating selection acting on *Pseudomonas*, such that it became specifically adapted to evolving *Variovorax*, while becoming maladapted to ancestral *Variovorax*.

**Figure 4.**
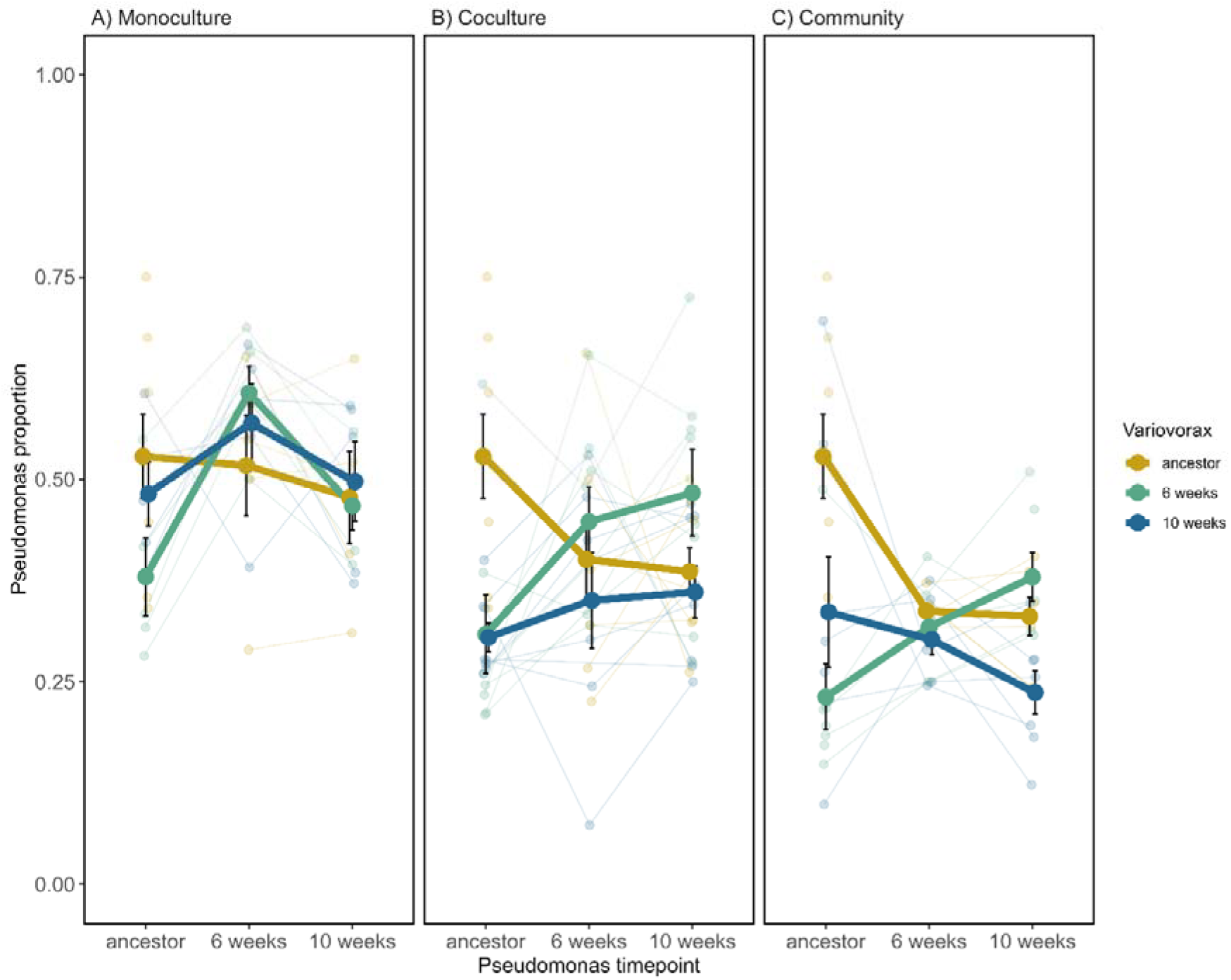
Relative proportion of Pseudomonas when cultured with Variovorax from different time-points. Pseudomonas and Variovorax evolved in (A) monoculture (each species in isolation), (B) in coculture (Pseudomonas and Variovorax) and in (C) community (Pseduomonas and Variovorax embedded with 3 other bacterial species). Patterns of coevolution present for (B) and (C) but not (A) observed as significant interaction between evolutionary time of Pseudomonas and Variovorax. Independent treatment replicates are small points, large points are mean proportion of Pseudomonas, bars represent ± SE. Thin lines are connecting the replicates that are tracked through time (except for the ancestor), thick lines represent the mean. Ancestor-ancestor combinations between the panels are the same 8 replicates presented to aid visualization.

To control for adaptation to abiotic conditions potentially being interpreted as coevolution, monoculture lines were evolved alongside the coculture and community treatments. We did not detect interactions that were suggestive of coevolution between *Pseudomonas* and *Variovorax* evolved in monoculture (GLMM: *Pseudomonas* time x *Variovorax* time: χ2_3_ = 4.02, p = 0.26). There was no indication of *Variovorax* adapting to abiotic conditions or that any abiotic adaptation affected its interaction with *Pseudomonas* (GLMM: *Variovorax* time χ2_2_ = 0.88, p = 0.64, Figure 3A). We found some evidence for abiotic adaptation in *Pseudomonas, which* affected interactions with *Variovorax* evolved in monoculture, with an increase in the proportion of *Pseudomonas* at week 6 (GLMM: *Pseudomonas* time χ2_2_ = 10.06, p = 0.007, Figure 4A), but not week 10.

### Community complexity did not affect coevolution of species

Contrary to our expectation that evolution in a multispecies community would weaken coevolution, we found no difference in coevolutionary dynamics between the coculture and community treatments (GLMM: *Pseudomonas* time x *Variovorax* time x complexity interaction χ^2^_3_ = 3.54, p = 0.32). *Pseudomonas* proportion was not differentially affected by community complexity over evolutionary time for each species (GLMM: *Pseudomonas* time x complexity interaction and *Variovorax* time x complexity interaction: χ^2^_2 pseudomonas_ = 4.01, p _pseudomonas_ = 0.14; χ^2^_2 variovorax_ = 4.21, p_variovorax_ = 0.12). However, selection pressures are clearly different between the two treatments. *Pseudomonas* evolving in community context displayed lower proportions compared to cocultures (Tukey HSD: estimate = 0.33, z-ratio = 5.07, p < 0.001), suggesting that *Pseudomonas* adaptation to *Variovorax* is weakened in a community context. This is driven by both an increase in *Variovorax* density (Tukey HSD: estimate = -0.08, t-ratio = -3.10, p = 0.003) and a decrease in *Pseudomonas* density (Tukey HSD: estimate = 0.06, t-ratio = 2.12, p = 0.037, Figure 5C).

**Figure 5.**
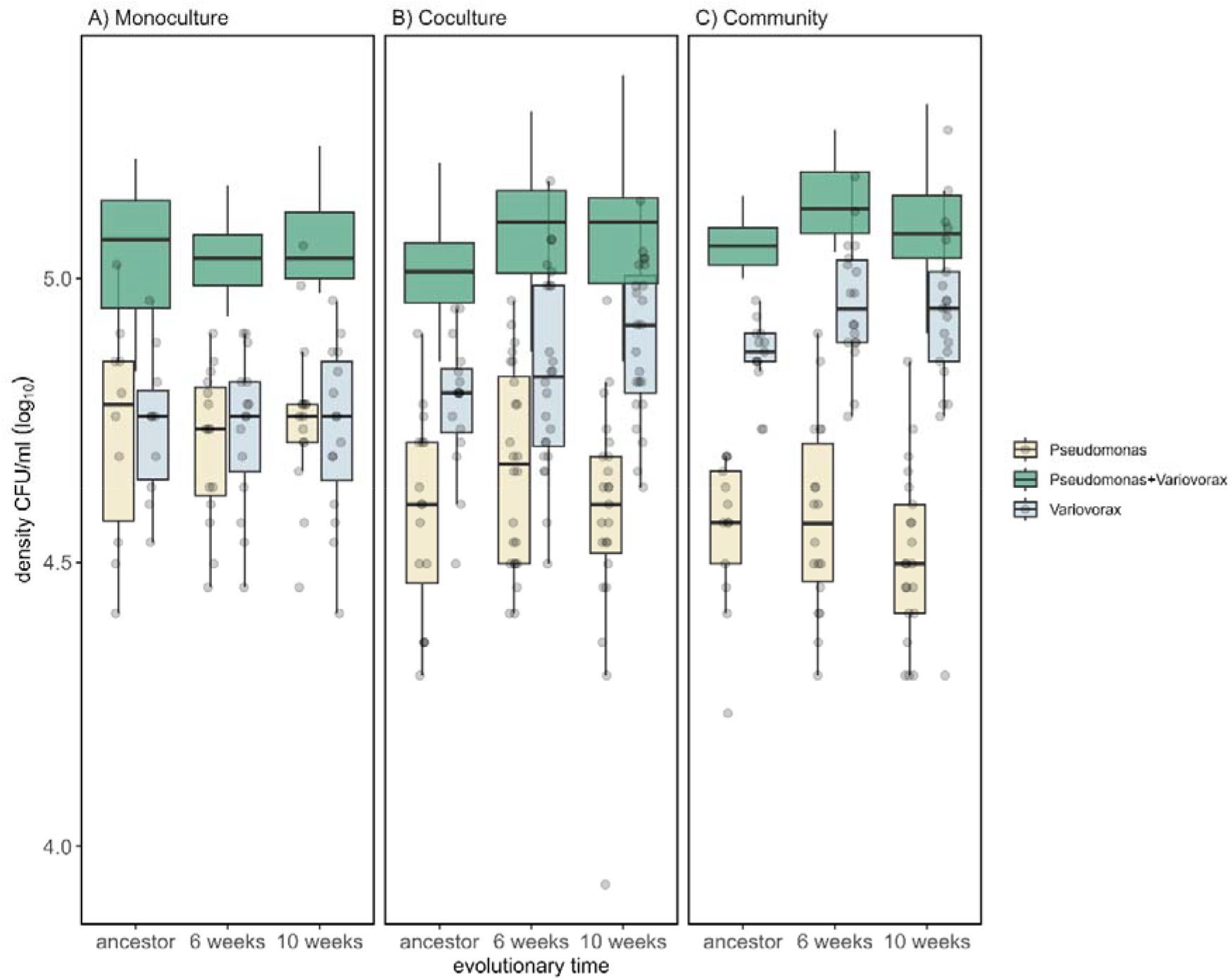
The density of individual species and their combined density when assembled as two-species cocultures (Variovorax and Pseudomonas) following evolution in (A) monoculture (each species evolved alone), (B) coculture (Pseudomonas and Variovorax only), and (C) in a community (with three other species). Population densities are plotted against evolutionary time. Boxplots showing medians, first and third quartiles, whiskers are 1.5 * interquartile range (IQR). Individual points represent individual treatment replicates.

### Changes in total density driven by Variovorax evolution

Total density (of *Pseudomonas* and *Variovorax* cocultures) could change over evolutionary time due to a change in species interactions (e.g. increased exploitation by *Variovorax* could result in a decline in total density). We observed an increase in total density as a result of an increase in exploitation of evolved *Variovorax*, and this effect was independent of evolutionary time and background (community complexity) (LMM: *Pseudomonas*_time_ x *Variovorax*_time_ x community complexity interaction: χ^2^_6_ = 5.11, p = 0.53, Figure 5). Total density was not differentially affected by community complexity over evolutionary time for each species (LMM: 2-way interaction for *Pseudomonas*_*time*_ x complexity and *Variovorax*_*time*_ x complexity: χ^2^_4 *pseudomonas*_ = 6.58, p _*pseudomonas*_ = 0.16; χ^2^_4 *variovorax*_ = 2.72, p _*variovorax*_ = 0.61). Total density was also not affected by an interaction between the two species evolutionary time (LMM: *Pseudomonas* x *Variovorax* time interaction: χ2_3_ = 5.90, p = 0.12). Only the evolutionary time of *Variovorax* had a significant effect on the density of the assembled co-cultures (LMM effect of *Variovorax* time: χ2_2_ = 7.53, p = 0.024). This is driven by an increase in *Variovorax* density at week 6 (Tukey HSD: estimate = - 0.054, t-ratio = -2.57, p = 0.047) leading to an increase in total density compared to the ancestor (Figure 5BC). Therefore, increased exploitation of *Pseudomonas* by *Variovorax* increased community density despite decreases in *Pseudomonas* density.

## Discussion

In this study, we sought to understand the effect of community complexity on exploitation-mediated pairwise coevolution in a soil microbial community. In our model system *Pseudomonas* (and some other members of the community) are exploited by *Variovorax* mediated by interactions over metabolites (Castledine et al., 2024). While we found evidence for antagonistic coevolution, community complexity did not significantly affect coevolutionary dynamics. This is despite the community context providing more species for *Variovorax* to exploit (therefore reducing selection on *Pseudomonas* specifically); and *Pseudomonas* experiencing competition from more species, therefore inhibiting population sizes and/or offering conflicting selection pressures. This suggests that reciprocal selection between *Pseudomonas* and *Variovorax* is sufficiently strong to buffer the effect of the altered selection pressures created by the community context.

While there are examples of within-trophic level studies of bacterial interactions showing how species interactions can lead to different evolutionary outcomes in both coculture and multispecies communities (Chang et al., 2020; Chen & Zhang, 2024; Pearl Mizrahi et al., 2023; Piccardi et al., 2024), to our knowledge this is the first study explicitly demonstrating within-trophic level antagonistic coevolution between bacteria. The mechanisms underlying any observed adaptations in this study are unknown. However, from previous work we know that *Variovorax* benefits from the presence and metabolic activity of *Pseudomonas* in our model (Castledine et al., 2024), therefore mechanisms of *Pseudomonas* resistance to *Variovorax* are likely related to alteration of metabolites, while *Variovorax* becomes more efficient at using metabolites produced by *Pseudomonas* or adapts to utilise altered metabolites. Analysing the exact nature of this interaction (and evolutionary mechanisms thereof), is made difficult owing to the complex nutrient medium.

In both pairwise cocultures and communities, *Variovorax* became more efficient at exploiting *Pseudomonas*, with *Variovorax* reaching higher densities relative to *Pseudomonas* through time. Exploitation of *Pseudomonas* by *Variovorax* showed an overall increase through time that is consistent with arms race dynamics (Gandon et al., 2008). In turn, *Pseudomonas* adapted by becoming more resistant to exploitation, albeit with a time lag, to 6-week evolved *Variovorax*. However, there was no observed increase overall in resistance of *Pseudomonas* against *Variovorax* over time, with the increase in resistance to week 6 *Variovorax* being accompanied by a decrease in resistance to ancestral *Variovorax*. This suggest*s* that fluctuating selection dynamics (FSD) acting on *Pseudomonas* played an important role in the coevolution within this system (Hall et al., 2011). Our knowledge of antagonistic coevolutionary dynamics in microbes is mostly based on bacteria-phage models, where selection pressures and the evolutionary potential are different to within-trophic coevolution (Buckling & Rainey, 2002; Gandon et al., 2008; Gómez & Buckling, 2011). Infection with lytic phage leading to cell death imposes strong selection on bacteria to adapt defences, whereas competitive interaction between bacteria – especially those involving interactions over metabolites - might be less specific, leading to weaker selection. FSD in host-pathogen systems becomes more prevalent due to increasing costs of higher infectivity/resistance for the phage and bacteria, respectively (Hall et al., 2011), resulting in selection on standing genetic variation. This might explain why most work to date, using a bacteria-phage model system reported coevolving partners both exhibiting either ARD or FSD. Our results suggest that this is not the case for within trophic-level coevolution, where coevolving partners show different coevolutionary dynamics. Further work is needed to explore the wider implications of such highly asymmetric coevolutionary dynamics and their impact on ecology and evolution.

A common criticism of laboratory studies of coevolution is that they are overly simplistic in their conditions and far removed from nature to offer insight into more complex evolutionary dynamics. Previous studies have found that even small increases in community complexity can significantly affect coevolution (Barraclough, 2015; Blazanin & Turner, 2021; Castledine, Sierocinski, et al., 2022; Manriquez et al., 2021). Our results instead find that pairwise coevolution can predict coevolution in community contexts which may be due to selection pressures being sufficiently strong. Similar cases of parallel evolutionary dynamics may be observed in wider contexts such as phage therapy, where bacteria experience strong selection to phage in patients and in closed laboratory conditions (Castledine, Padfield, et al., 2022). As *Pseudomonas* is the species *Variovorax* derives the strongest fitness benefit from, interaction intensities and reciprocal selection may have been sufficiently strong for coevolution despite co-occurring community members. Mutation supply rates may have also been non-significantly affected by other community members as *Variovorax* can generally exploit at least two other community members (other than *Pseudomonas*) which may allow it to maintain sufficient mutation rates for coevolution (Castledine et al., 2024; Gandon & Michalakis, 2002). Wider work analysing coevolutionary interactions within trophic levels, including between cross-feeding mutualists and exploitative interactions, will give insights into how coevolution occurs in natural community contexts. The observed negative effect of community complexity on evolved *Pseudomonas* densities might be explained by additional selection pressure arising from competition with other community members. *Variovorax* benefits from most species in the community, while *Pseudomonas* competes against them leading to stronger selection on *Pseudomonas* in the community treatment. This potentially leads to trade-offs between adaptation to *Variovorax* and other community members.

In this study, we explicitly demonstrate antagonistic coevolution between naturally co-occurring bacteria. This antagonistic coevolution leads to increased exploitation but not increased resistance over the experimental time and shows an important role of both ARD and FSD. Furthermore, we show that pairwise coevolution can be robust in the face of community complexity. Coevolutionary dynamics did not change significantly with increased complexity, despite finding differences in densities of species suggesting a change in selective pressures. Understanding the interplay between coevolution and biotic complexity is crucial not only for advancing evolutionary theory but also for applications in medicine, biotechnology, and ecology, where microbial coevolution can influence antibiotic resistance, pathogen evolution, and microbiome stability. This work contributes to our understanding of within-trophic level coevolution, with implications for both natural and engineered microbial communities.

